# Reduced DPP4 Binding Confers Resistance to Soluble DPP4 While Preserving MERS-CoV Entry into Cells Expressing High Levels of DPP4

**DOI:** 10.64898/2026.09.14.751648

**Authors:** Nianzhen Chen, Stefan Pöhlmann, Markus Hoffmann

**Affiliations:** Infection Biology Unit, German Primate Center –Leibniz Institute for Primate Research, 37077 Göttingen, Germany; Faculty of Biology and Psychology, Georg-August University Göttingen, 37073 Göttingen, Germany; Institute of Molecular Virology, Ulm University Medical Center, Ulm, Germany; German Center for Infection Research (DZIF), associated partner site Göttingen, Göttingen, Germany

**Author notes:** Address correspondence to Markus Hoffmann,; Stefan Pöhlmann.

**Keywords:** Middle East Respiratory Syndrome, spike, DPP4, soluble DPP4

## Abstract

The Middle East respiratory syndrome coronavirus uses DPP4/CD26 as receptor for cell entry. Soluble recombinant DPP4 can block MERS-CoV infection in cell culture and in animal models and might hold promise as antiviral. Furthermore, endogenous soluble DPP4 in plasma might reduce MERS-CoV dissemination in infected individuals. Therefore, we addressed whether and how MERS-CoV can acquire resistance against soluble DPP4 (sDPP4), employing a vesicular stomatitis virus (VSV) encoding the MERS-CoV spike (S) protein (VSV-MERS-S). Passaging of VSV-MERS-S in the presence of sDPP4 selected for viral variants with mutations L507I and L507H in the receptor binding domain (RBD) of the S protein. These mutations reduced sDPP4 binding, were compatible with robust entry into cell lines expressing high levels of DPP4 and conferred sDPP4 resistance. In contrast, entry into cell lines expressing low levels of DPP4 was reduced. Furthermore, polymorphisms L507F/R/P, which were detected in MERS-CoV sequences from patients, exhibited an even more pronounced phenotype, with L507R and L507P conferring complete sDPP4 resistance. Collectively, our results show that naturally occurring polymorphisms in the MERS-CoV S protein can confer sDPP4 resistance by reducing sDPP4 binding but might still be compatible with robust viral spread in cells and tissues expressing high levels of DPP4.

**IMPORTANCE:** DPP4 is the cellular receptor used by MERS-CoV for entry into target cells. A soluble form of DPP4 (sDPP4) blocks MERS-CoV infection and has been proposed as a potential antiviral. Here, using surrogate systems, we show that MERS-CoV can acquire resistance to sDPP4 through spike protein mutations that reduce DPP4 binding. These mutations remain compatible with efficient entry into cells expressing high levels of DPP4 and were also identified in patient-derived MERS-CoV. Our findings suggest that MERS-CoV can balance receptor engagement with resistance to a soluble receptor decoy, with implications for antiviral strategies targeting virus-receptor interactions.

---

The Middle East respiratory syndrome coronavirus (MERS-CoV) is a betacoronavirus that naturally infects dromedary camels, in which infection is typically asymptomatic or associated with mild rhinitis (1). Infected camels can transmit the virus to humans, where infection may cause severe respiratory disease and renal failure with a case-fatality rate exceeding 30%, particularly in individuals with underlying comorbidities (1). Human-to-human transmission of MERS-CoV is currently inefficient and occurs primarily in healthcare settings. However, there is concern that the virus could adapt for more efficient replication in the upper respiratory tract, thereby increasing its transmissibility.

MERS-CoV was first identified in humans in 2012 (2), and sporadic human infections continue to occur (3). A marked decline in reported cases was observed during the COVID-19 pandemic (3) but it remains unclear whether this resulted from public health interventions implemented to control SARS-CoV-2 or cross-protective immunity induced by SARS-CoV-2 infection or vaccination, with a recent study reporting that post-pandemic sera did not show increased neutralization activity against MERS-CoV (4). Although most cases have occurred in the Middle East, infected travelers have repeatedly introduced MERS-CoV into other countries, resulting in several outbreaks (3). The largest outbreak outside the Arabian Peninsula occurred in South Korea in 2015, where a single infected traveler initiated a hospital-associated outbreak involving 186 laboratory-confirmed cases and 38 deaths (5). Collectively, these observations highlight the continued pandemic potential of MERS-CoV, which the World Health Organization (WHO) has designated a priority pathogen for epidemic preparedness.

At present, there are no licensed antivirals or vaccines to combat MERS-CoV infection. Coronavirus spike proteins, including the MERS-CoV spike protein (MERS-S), facilitate viral entry into host cells, are the key target of the neutralizing antibody response and constitute a potential point of attack for novel antivirals (6–8). For entry into cells, the receptor binding domain (RBD) of the S protein binds to cell surface protein DPP4/CD26, a serine exopeptidase that processes diverse substrates and is involved in multiple biological processes, including energy metabolism and immune regulation (7, 8). DPP4 is inserted into the plasma membrane via a transmembrane domain but can be shed into the extracellular space upon cleavage of its membrane proximal region by host cell proteases (9, 10) and high levels of soluble DPP4 (sDPP4) are found in human plasma (11, 12). Previous studies reported that recombinant sDPP4 can inhibit MERS-CoV entry into cultured cells (8, 13). Furthermore, reduced susceptibility to MERS-CoV infection in mice homozygous for human DPP4, relative to heterozygous animals, correlated with higher circulating levels of human sDPP4 (13). These findings suggest that sDPP4 can exert antiviral activity in cell culture and in the infected host. Whether the concentration of endogenous sDPP4 in the plasma of patients is sufficient to inhibit MERS-CoV infection has been unclear, with a recent study providing indirect evidence for such a scenario (14). Collectively, recombinant sDPP4 can exert antiviral activity when supplied at sufficient levels and may merit the development as novel antiviral while further investigation is required to determine whether endogenous sDPP4 can interfere with MERS-CoV infection.

Here, we investigated whether and how MERS-CoV can acquire resistance to sDPP4. Employing chimeric vesicular stomatitis virus (VSV) encoding MERS-S (VSV-MERS-S) and S protein bearing pseudotypes as model systems, we show that cell culture selected and naturally occurring mutations at position L507 can reduce sDPP4 binding and susceptibility to sDPP4 inhibition but are compatible with entry into cell lines expressing high levels of DPP4. Our findings indicate that MERS-CoV can modulate receptor binding to evade inhibition by a soluble receptor decoy, highlighting potential challenges for antiviral strategies targeting virus-receptor interactions.

## RESULTS

### Selection of MERS-S mutations that allow for viral spread in the presence of sDPP4

In order to investigate how MERS-CoV could develop resistance against sDPP4, we employed VSV-MERS-S, which encodes the spike of MERS-CoV 2c EMC/2012 isolate (wildtype, WT). This chimeric virus enters cells in a MERS-S-dependent fashion and is replication competent, allowing for selection of sDPP4-resistant mutants. For mutant selection, Caco-2 cells were inoculated with VSV-MERS-S in the presence of serially diluted sDPP4 and incubated for 24 to 48 h (Fig. 1A). Subsequently, the cells showing signs of infection in the presence of the highest concentration of sDPP4 were identified by fluorescence microscopy, supernatants collected, diluted and transferred to fresh target cells in the presence of serially diluted sDPP4 (Fig. 1A). This procedure was carried out with technical triplicates and repeated 10 times. Subsequent sequence analysis revealed that a fraction of the viral inoculum contained mutation L697F in MERS-S, which likely constitutes a pre-existing, cell-culture-derived variant that arose during virus rescue or initial propagation. After 5 and 10 passages in the presence of sDPP4, MERS-S mutation L507H was detected in one of the three replicates (Fig. 1B). In another replicate, mutations L507I and T533P were found after 10 passages while no mutations were found in the remaining replicate (Fig. 1B). These results suggest that selective pressure exerted by sDPP4 might select for mutations at position L507 and potentially also at position T533, both located in the RBD of MERS-S protein (Fig. 1C).

**FIG 1.**
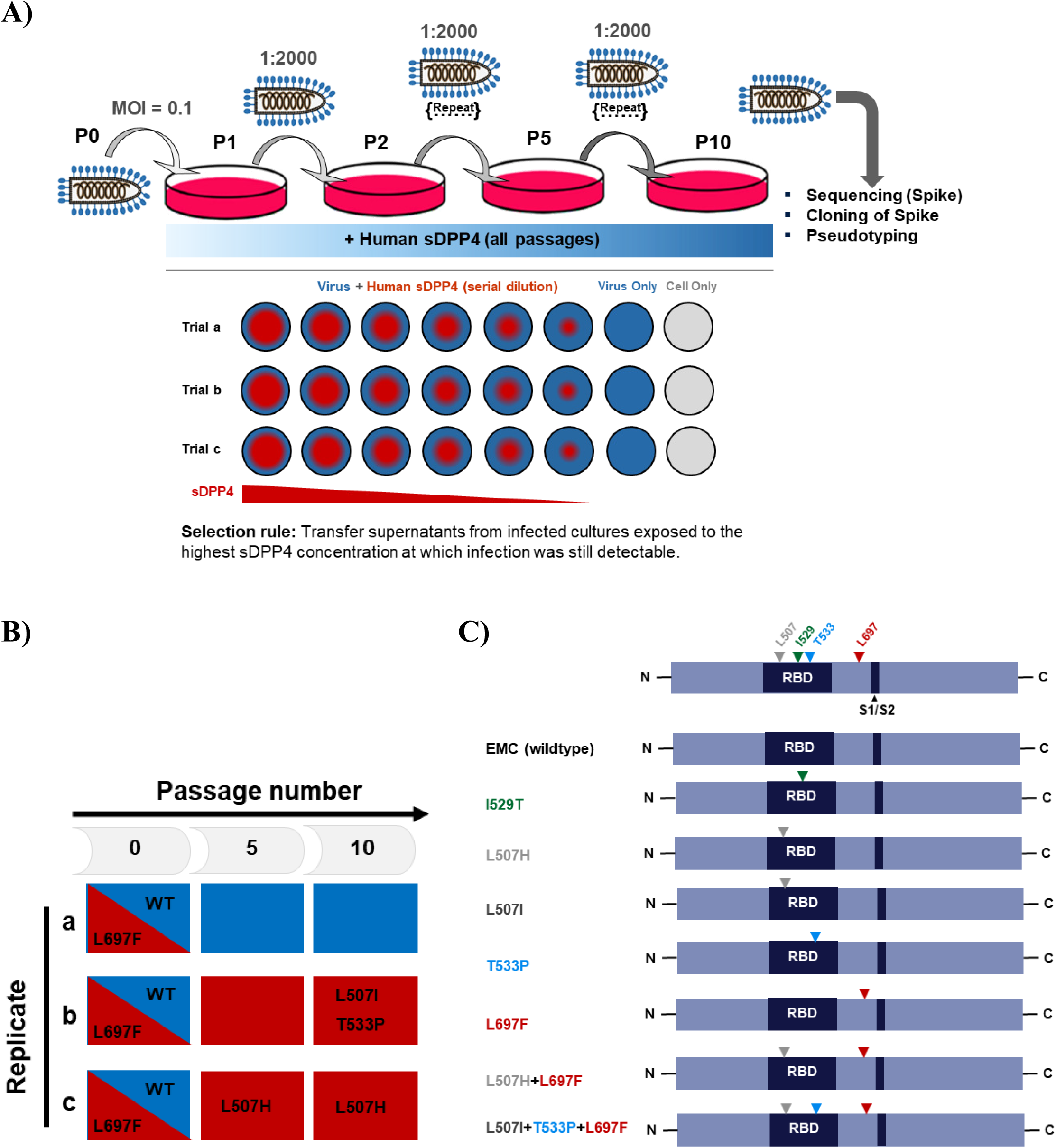
Selection for MERS-S mutants resistant against sDPP4. A) Schematic representation of the in vitro evolution of VSV-MERS-S under sDPP4 selection pressure. Caco-2 cells were inoculated with VSV-MERS-S (passage 0, P0), and sDPP4 at the indicated serial dilutions was added after virus adsorption. At 24–48 h postinfection, supernatants were collected from the well containing the highest sDPP4 concentration that still permitted efficient infection. Supernatants were clarified to remove cellular debris, diluted 1:2,000, and used to inoculate fresh Caco-2 cells. sDPP4 was then added again at serial dilutions to initiate the next passage. This selection scheme was continued for 10 passages (P1–P10), and the resulting supernatants were subjected to downstream phenotypic and genotypic analyses. B) Amino acid substitutions identified in MERS-S across three independent replicates (a–c) at passages 0 (P0), 5 (P5), and 10 (P10). Blue indicates wild-type (WT, MERS-S EMC) residues, and red indicates substituted residues. At P0, L697F was detected as a mixed population. L507H, L507I and T533P were detected at later passages in a replicate-dependent manner. C) Mutations analyzed in the present study and their localization within the MERS-S protein. Triangles mark the positions of amino acid substitutions identified in this study. RBD, receptor-binding domain; S1/S2, motif for proteolytic processing.

### L507H and L507I reduce entry into cell lines that express low levels of DPP4 but are compatible with robust entry into cell lines expressing high levels of DPP4

To elucidate whether the MERS-S mutations detected upon passaging of VSV-MERS-S in the presence of sDPP4 indeed confer resistance to sDPP4, the single mutations or combinations thereof were introduced into MERS-CoV S protein (Fig. 1C) and their impact on cell entry studied with single-cycle VSV reporter particles pseudotyped with the S protein. MERS-S with mutation I529T was included as control, since I529T was previously shown to reduce DPP4 binding (15, 16). Immunoblot analyses revealed that all S protein mutants were comparably incorporated into VSV particles although incorporation of I529T and the triple mutant L507I, T533P and L697F appeared to be moderately reduced as compared to other S proteins tested, although effects were not statistically significant (Fig. 2A).

**FIG. 2.**
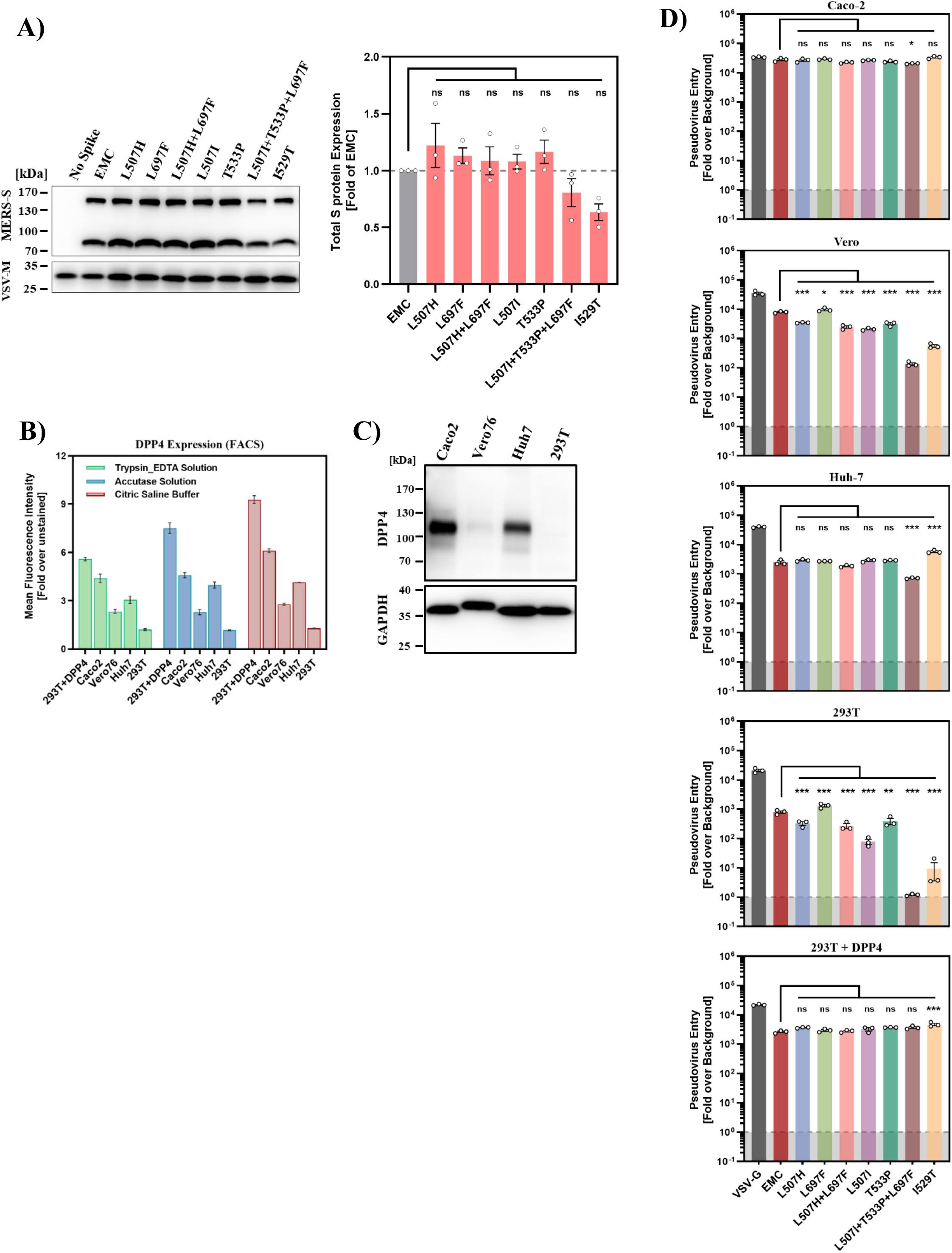
Mutations arising under sDPP4 selection (L507H, L507I) reduce entry into cell lines expressing low levels of DPP4 but are compatible with efficient entry into cell lines expressing high levels of DPP4. A) S protein incorporation into particles and S1/S2 cleavage. Pseudotyped viral particles bearing MERS-S with a C-terminal V5-tag were analyzed by immunoblotting to assess MERS-S particle incorporation and proteolytic processing. The uncleaved S protein and the S2 subunit were detected using an anti-V5 antibody. Detection of VSV-M protein served as loading control. A representative blot from three independent experiments is shown (left panel). Quantification of S protein incorporation into particles is shown in the right panel. Incorporation was determined by normalizing total S protein band intensities to the corresponding VSV-M signal for each sample and then expressing values relative to EMC S (set to 1). Data represent mean ± SEM from three independent biological replicates. Statistical analysis was performed using one-way analysis of variance (ANOVA) with Dunnett’s posttest (ns, not significant). B) DPP4 surface expression. Surface expression of DPP4 was quantified by flow cytometry in Caco-2, Vero, Huh7, 293T and 293T-DPP4 cells using an anti-DPP4 antibody and an Alexa Fluor 488–conjugated secondary antibody (left panel). Surface DPP4 levels are shown as mean fluorescence intensity (MFI) normalized to the unstained control (set to 1). Data represent mean ± SEM from three independent experiments. Similar results were obtained regardless of the method used for cell detachment. C) Total DPP4 expression. Whole-cell lysates from the indicated cell lines were immunoblotted for DPP4, with GAPDH serving as a loading control. A representative blot is shown and results were confirmed in two independent experiments. D) Entry into cell lines. The indicated cell lines were inoculated with pseudotyped particles bearing the S proteins of the indicated MERS-CoV variants, wild-type MERS-CoV S (EMC) or VSV-G as a positive control. Transduction efficiency was quantified by measuring virus-encoded luciferase activity in cell lysates at 16–20 h post transduction. Presented are the average (mean) ± SEM data from three biological replicates (each conducted with technical quadruplicates). Statistical analysis was determined by one-way analysis of variance (ANOVA) with Dunnett’s posttest (ns, not significant; *, p ≤ 0.05; **, p ≤ 0.01; ***, p ≤ 0.001).

For analysis of MERS-S-driven entry, Caco-2, Huh-7, Vero, 293T and 293T cells transiently expressing DPP4 (293T + DPP4) were used as targets. These cell lines differ in DPP4 expression as determined by fluorescence-activated cell sorting (FACS) upon detachment of cells from culture flasks using trypsin and EDTA, accutase or citric saline buffer (Fig.2B). Although maximum fluorescence differed depending on the mode of detachment, consistent results were obtained when examining relative DPP4 expressing levels. Highest levels were measured for 293T + DPP4 cells, as expected, followed by Caco-2, Huh-7, Vero and 293T cells (Fig. 2B). Similar results were obtained upon immunoblot analysis (Fig. 2C), confirming that the target cell lines chosen differed markedly in DPP4 expression.

Analysis of cell entry driven by the MERS-S mutants analyzed revealed a clear impact of DPP4 levels on entry efficiency. All mutant S proteins facilitated entry into 293T-DPP4 and Caco-2 cells, the cell lines with highest DPP4 expression, with comparable efficiency as the WT MERS-S protein (Fig. 2D). Despite the high overall cell entry efficiency in both cell lines, particles bearing the L507I/T533P/L674F mutant showed a slight but statistically significant increase in entry into 293T + DPP4 cells, whereas entry into Caco-2 cells was slightly but significantly reduced compared with MERS-S WT.. Similar results were obtained for Huh-7 cells, which expressed slightly reduced DPP4 levels as compared to Caco-2, although the reduction in entry driven by mutant L507I+T533P+L697F was more pronounced relative to Caco-2 cells (Fig. 2D). A profound reduction of cell entry driven by mutant MERS-S proteins was observed for Vero cells, which expressed markedly less DPP4 than Huh-7 cells. The only exception was mutant L697F, which facilitated only slightly, but statistically significantly, reduced entry into this cell line compared to MERS-S WT (Fig. 2D). Finally, a phenotype even more pronounced than that observed for Vero cells was obtained for 293T cells, which expressed the lowest levels of DPP4. Most strikingly, entry into this cell line driven by S protein mutants L507I+T533P+L697F and I529T was close to or within the background level (Fig. 2D). Thus, the MERS-S mutations selected during passaging in the presence of sDPP4 remained compatible with robust entry into cell lines expressing high, but not low, levels of DPP4.

### L507H und L507I reduce DPP4 binding and susceptibility to inhibition by sDPP4

We next investigated whether the S protein mutations analyzed impacted binding to DPP4. For this, binding of sDPP4 to S protein expressing cells was analyzed. FACS analysis revealed efficient cell-surface expression of all S proteins, although expression of mutants T533P and L507I+T533P+L697F was slightly reduced. (Fig. 3A). Mutation I529T markedly reduced sDPP4 binding, as expected (Fig. 3A, Fig. S1A). Similarly, mutation L507H and, particularly, L507I significantly reduced binding to sDPP4, in agreement with L507 being located close to the S protein residues contacting DPP4 (Fig. S2), while mutations T533P and L697F had no appreciable effect (Fig. 3A, Fig. S1A). Finally, combining mutation L507H with L697F did not result in further reduced sDPP4 binding as compared to L507H alone, while the combination of L507I+T533P+L697F markedly diminished sDPP4 binding as compared to the single mutations. Thus, the mutation L507H and particularly L507I reduced sDPP4 binding.

**FIG. 3.**
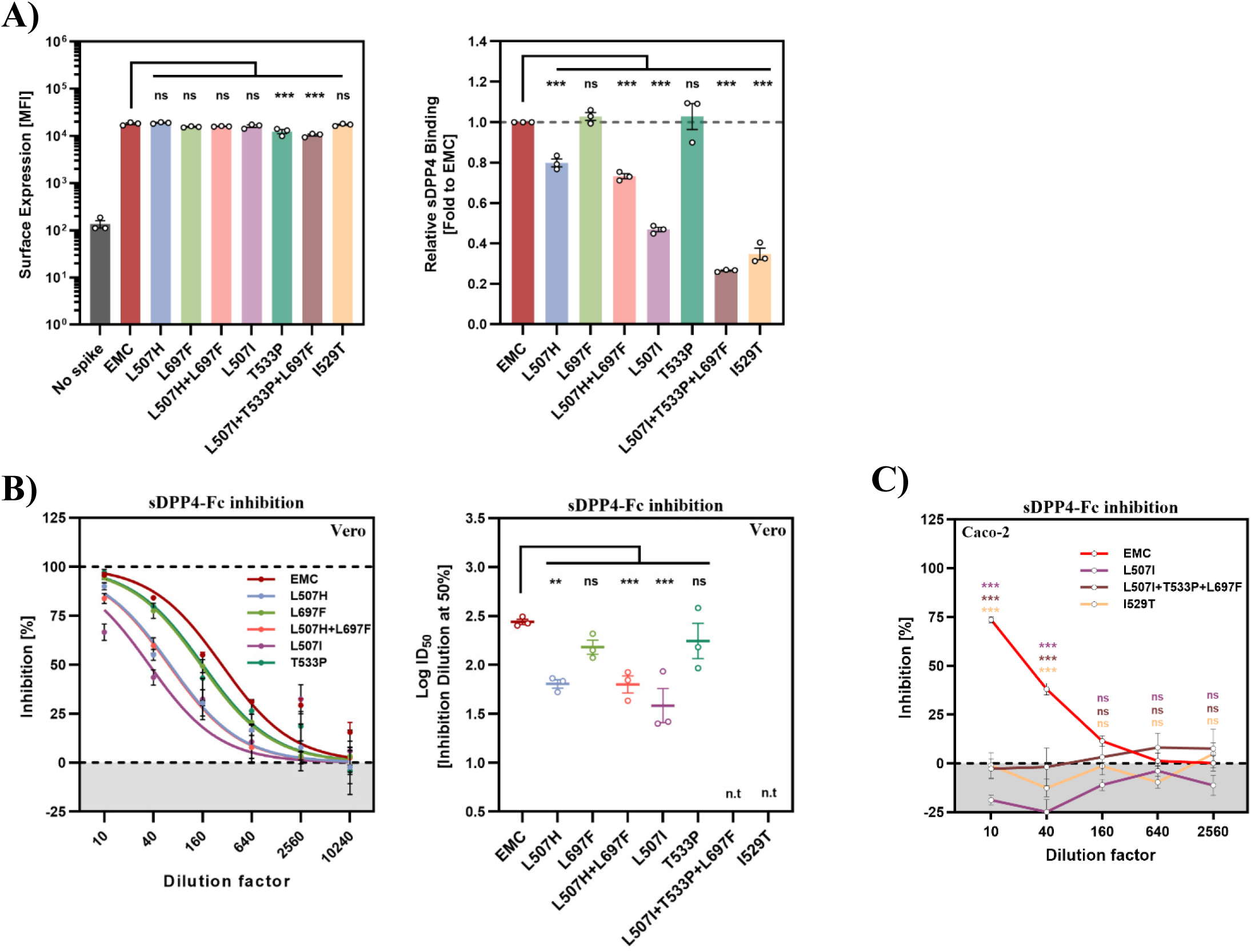
L507H and L507I reduce sDPP4 binding and susceptibility to inhibition by sDPP4. A) Surface expression and sDPP4 binding of MERS-S mutants. 293T cells transiently expressing the indicated S proteins (or empty vector control) were analyzed by flow cytometry using anti-MERS-S antibody followed by an Alexa Fluor 488-conjugated secondary antibody (left panel). In parallel, cells were incubated with sDPP4 and an Alexa Fluor 488-conjugated anti-human antibody to evaluate DPP4 binding. Relative DPP4 binding (right panel) represents DPP4 binding normalized to S protein surface expression with results obtained for EMC S protein set as 1. Data represent means ± SEM from three independent experiments. Statistical significance was determined using a one-way analysis of variance (ANOVA) with Dunnett’s posttest (ns, not significant; *, P ≤ 0.05; **, P ≤ 0.01; ***, P ≤ 0.001). B) Inhibition of Vero cell entry by sDPP4. VSV pseudotyped with the indicated S proteins were pre-incubated with serially diluted sDPP4 for 30 min at 37°C and then inoculated onto Vero cells. Spike-mediated entry was quantified 16–20 h postinfection by measuring virus-encoded luciferase activity in cell lysates. Entry of particles in the absence of sDPP4 was set as 0% inhibition. Inhibition curves obtained with increasing sDPP4 dilutions are shown in the left panel. The 50% inhibitory dilution (ID₅₀) was calculated by nonlinear regression and is shown in the right panel. Data represent the mean ± SEM from three independent experiments performed in technical quadruplicate. Statistical significance was determined by one-way analysis of variance (ANOVA) with Dunnett’s posttest (ns, not significant; *, P ≤ 0.05; **, P ≤ 0.01; ***, P ≤ 0.001). n.t, not tested. C) The experiment was carried out as described for panel B but Caco-2 cells were used as targets. Entry inhibition by sDPP4 is shown. Presented are the average (mean) data ± SEM from three biological replicates (each performed with technical quadruplicates). Statistical significance relative to MERS-S EMC was assessed by two-way ANOVA with Dunnett’s posttest. ns, not significant; *, P ≤ 0.05; **, P ≤ 0.01; ***, P ≤ 0.001.

Next, we analyzed the impact of the S protein mutations on susceptibility to entry inhibition by sDPP4, comparing Vero cells, which express low levels of DPP4, and Caco-2 cells, which express high levels of DPP4. The amount of inoculum was adjusted to allow for comparable entry of all pseudotyped particles studied in the absence of sDPP4, except for S protein mutants L507I+T533P+L697F and I529T, which failed to facilitate appreciable entry into Vero cells (Fig. S1B) but mediated robust entry into Caco-2 cells (Fig. S1C). sDPP4 efficiently blocked S protein-driven entry into Vero cells and mutation L507H and particularly L507I conferred resistance to inhibition by sDPP4 while L697F and T533P did not (Fig. 3B). Inhibition of Vero cell entry of mutants L507I+T533P+L697F and I529T by sDPP4 could not be analyzed due to inefficient entry in the absence of sDPP4 (Fig. S1B). In contrast, Caco-2 cell entry driven by S protein mutants L507I, L507I+T533P+L697F and I529T was robust and comparable to that measured for MERS-S WT (Fig. S1C). sDPP4 inhibited Caco-2 cell entry driven by the WT S protein with reduced efficiency as compared to Vero cell entry and the three mutants studied were completely resistant (Fig. 3C). These finding suggest that mutations L507H and L507I reduce sDPP4 binding and thereby confer resistance against sDPP4. In the context of target cells expressing low levels of DPP4 this comes at the cost of reduced entry efficiency in the absence of sDPP4 while entry into cells expressing high levels of DPP4 proceeds unabated.

### Naturally occurring mutations L507F and L507R confer resistance against sDPP4

We finally investigated whether residue 507 is polymorphic in MERS-CoV isolates and whether naturally occurring mutations at this residue impact sDPP4 binding and susceptibility to inhibition by sDPP4. Analysis of MERS-S sequences deposited in Genbank revealed that mutations L507F, L507R and L507P occur naturally (17, 18). These mutations were compatible with robust S protein expression and virion incorporation with the exception of L507R, which reduced S protein levels on VSV particles (Fig. 4A-B). The mutations were also compatible with robust entry into Caco-2 and Huh-7 cells, which express medium to high levels of DPP4, although entry driven while L507F slightly decreased entry efficiency. In contrast, L507F, L507R and, particularly, L507P markedly reduced entry into Vero cells, which express low levels of DPP4 (Fig. 4C). Further, mutation L507P dramatically reduced sDPP4 binding while L507F and L507R exerted a less severe effect comparable to that measured for mutations L507H and L507I (Fig. 4D, Fig. S1D). Finally, both L507H and L507I (selected in cell culture) and L507F and L507R (occurring naturally) markedly decreased susceptibility to inhibition by sDPP4, with L507R conferring complete resistance (Fig. 4E). The impact of mutation L507P could not be investigated due to its profound negative impact on Vero cell entry (Fig. 4C).

**FIG. 4.**
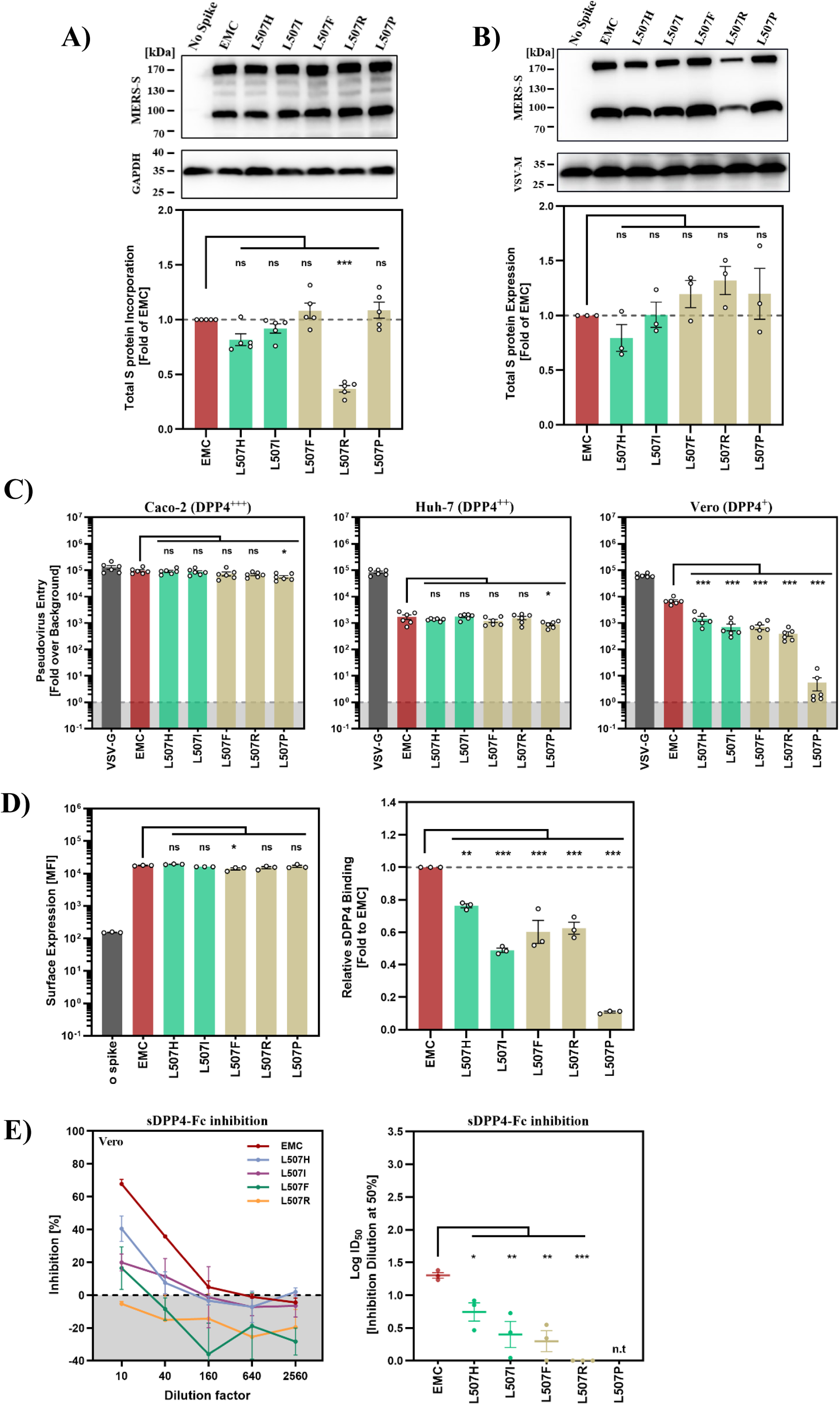
Naturally occurring mutations L507F and L507R confer sDPP4 resistance. A) Expression of S protein mutants. 293T cells were transfected with plasmids encoding the indicated C-terminally V5-tagged MERS-S proteins or an empty vector and subsequently infected with a replication-deficient VSV (VSV*ΔG-FLuc) to generate pseudotyped particles. Lysates from particle producing cells were analyzed by SDS-PAGE and immunoblotting using an anti-V5 antibody; GAPDH served as a loading control (top panel). Band intensities were quantified using ImageJ. Total S protein signals (S0+S2) were normalized to GAPDH (top left; n = 3) and expressed relative to EMC (set to 1). Data are mean ± SEM. Statistical significance was assessed by one-way ANOVA with Dunnett’s posttest (ns, not significant). B) Particle incorporation of S protein mutants. The particles generated as described in panel A were concentrated through a sucrose cushion and analyzed by immunoblotting using anti-V5 antibody for S protein detection (top panel). VSV-M served as a loading control. Band intensities were quantified using ImageJ. Total S protein signals (S0+S2) were normalized to VSV-M (top right; n = 5) and expressed relative to EMC (set to 1). Data are mean ± SEM. Statistical significance was assessed by one-way ANOVA with Dunnett’s posttest (ns, not significant). C) Cell line entry. Caco-2, Huh-7, and Vero cells were transduced with VSV pseudotyped particles bearing the indicated MERS-S mutants or MERS-S EMC. Particles bearing VSV-G served as a positive control. S protein-driven cell entry was analyzed at 16–18 h post inoculation by measuring luciferase activity in cell lysates and normalized against particles bearing no viral glycoprotein (set as 1). Data represent mean ± SEM from six independent biological experiments performed in technical quadruplicate. Statistical significance was determined by one-way analysis of variance (ANOVA) with Dunnett’s posttest (ns, p > 0.05; *, p ≤ 0.05; **, p ≤ 0.01; ***, p ≤ 0.001). D) sDPP4 binding. 293T cells expressing the indicated S proteins were analyzed by flow cytometry for both S protein surface expression (left panel) and sDPP4 binding (right panel). Relative sDPP4 binding represents sDPP4 binding normalized to S protein surface expression, with MERS-S WT set as 1. Data represent means ± SEM from three independent experiments. Statistical significance was determined using one-way analysis of variance (ANOVA) with Dunnett’s posttest (ns, not significant; *, P ≤ 0.05; **, P ≤ 0.01; ***, P ≤ 0.001). E) Entry inhibition by sDPP4. Pseudotyped particles bearing the indicated S proteins were incubated with serially diluted sDPP4 and subsequently used to infect Vero cells. S protein-dependent entry was quantified at 16–20 h postinfection by measuring luciferase activity in cell lysates and expressed relative to particles incubated in the absence sDPP4 (set to 0% inhibition). The dose-dependent inhibition curves (left panel) are shown. The 50% inhibitory dilution (ID₅₀) was determined by nonlinear regression analysis and is summarized on the right. Shown are the average (mean) data obtained from three biological replicates, each performed with four technical replicates. Statistical significance was determined one-way analysis of variance (ANOVA) with Dunnett’s posttest (ns, not significant; *, P ≤ 0.05; **, P ≤ 0.01; ***, P ≤ 0.001). n.t, not tested. Red, MERS-S EMC; green, S proteins from naturally occurring MERS-CoV variants, beige, spike mutants selected in cell culture.

## DISCUSSION

Endogenous sDPP4 is present at high concentrations in the blood, where it may interfere with the systemic spread of MERS-CoV (11, 19–22). Furthermore, recombinant sDPP4 has been shown to block MERS-CoV infection in cell culture and may hold promise as an antiviral agent (13, 23). However, it largely unclear whether and how MERS-CoV can acquire resistance to sDPP4. Our previous work showed that mutation Q1020R in the viral S protein reduces sensitivity to inhibition by sDPP4 but the mutation did not diminish DPP4 binding and it remained unknown how it conferred sDPP4 resistance (14). Here, using cell culture model systems, we provide evidence that spike mutations at residue L507, which emerged during cell culture passage of VSV-MERS-S and reduced sDPP4 binding, can confer resistance to sDPP4 without compromising viral entry into target cells expressing high levels of DPP4. Furthermore, we show that mutations at position L507 that were detected in MERS-CoV-infected patients recapitulate this cell culture phenotype. These findings suggest that MERS-CoV can tolerate mutations that reduce DPP4 affinity and confer resistance to sDPP4 without compromising its ability to infect target cells in the respiratory tract, which express high levels of DPP4.

Receptor decoys such as sDPP4 have been investigated for activity against several viruses. For example, a soluble form of CD4, the receptor for human immunodeficiency virus (HIV), was found to exhibit antiviral activity in cell culture (24–26). Fusion of CD4 to immunoglobulin Fc generated multivalent CD4 immunoadhesins with improved antiviral properties (27, 28), and an advanced version, the tetravalent CD4-IgG2 fusion protein PRO 542, reduced HIV-1 viral loads in infected patients (29–31). In the context of SARS-CoV-1 and SARS-CoV-2 infection, soluble forms of the ACE2 receptor have also been shown to exert antiviral activity in cultured cells and organoids (32, 33). The inhibition of infection was enhanced by ACE2 multimerization (34, 35) and by optimizing ACE2 affinity for the viral S protein (35), and protocols for administration via the respiratory tract have been established and safety was demonstrated (36, 37). However, clinical efficacy of systemically or topically applied soluble ACE2 in the context of coronavirus infection remains to be demonstrated. In the context of MERS, previous studies found that sDPP4 blocks infection in cell culture (13, 23). Moreover, elevated levels of sDPP4 are thought to contribute to the reduced susceptibility to MERS-CoV infection observed in transgenic mice homozygous for human DPP4 compared with heterozygous animals (23). Collectively, several lines of evidence suggest that recombinant sDPP4 and related receptor decoys hold promise as antiviral agents.

It is conceivable that not only exogenous but also endogenous sDPP4 inhibits MERS-CoV infection. Soluble DPP4 is present at high concentrations in several body fluids, including semen, urine, and serum/plasma (11, 12), due to shedding of DPP4 from the cell surface. Shedding results from cleavage of the DPP4 ectodomain by matrix metalloproteinases (MMPs) (10) or the serine protease KLK5 (9), with the cleavage site being located close to the transmembrane domain (38). Initially, cells of the kidney and hepatobiliary system were thought to be the principal sources of sDPP4, but more recent evidence suggests that T cells are major producers (12). One study reported that circulating sDPP4 levels are relatively low in the Saudi Arabian population and proposed that this may contribute to the high incidence of MERS in Saudi Arabia (39). In contrast, cell culture experiments indicated that physiological plasma concentrations of sDPP4 are below that needed to significantly inhibit MERS-CoV infection (13). Our previous work provided indirect evidence that plasma sDPP4 may moderately inhibit MERS-CoV entry into target cells (14). We further demonstrated that an S protein polymorphism present in Arabian MERS-CoV strains, Q1020R, reduces sensitivity to inhibition by sDPP4 compared with African strains (14), which may contribute, at least in part, to the absence of documented severe MERS cases in Africa despite widespread circulation of MERS-CoV in African camel populations.

The finding that recombinant sDPP4 exerts robust antiviral activity and the evidence suggesting that endogenous sDPP4 might be a natural defense against MERS-CoV infection highlight that it is important to determine whether the virus can acquire resistance against sDPP4. It can be speculated that sDPP4 and related receptor decoys might be associated with a higher barrier against resistance development as compared, for instance, to neutralizing antibodies targeting the receptor binding domain (RBD) in the viral S protein, since single amino acid exchanges can destroy the epitope of an antibody but in many cases will have little impact on receptor binding (40, 41). In agreement with this scenario, SARS-CoV-2 acquired a large number of mutations in the RBD to evade neutralizing antibodies but maintained efficient ACE2 usage throughout its evolution (42). In contrast, the present study shows that mutation of residue L507, acquired during passaging of a MERS-CoV surrogate model in the presence of rising amounts of sDPP4 was sufficient to confer sDPP4 resistance.

Residue L507 is located in the receptor binding motif (RBM), an exposed subdomain of the RBD that contacts DPP4 (7, 43). The residue itself does not make atomic contacts with DPP4 but, based on in silico predictions, is nevertheless important for DPP4 binding (44). Therefore, it is not surprising that mutations at L507 can decrease DPP4 binding. However, it is surprising that such mutations are selected for by sDPP4. Our data indicate that the reduced sDPP4 affinity associated with mutations L507F, L507R and L507P results in reduced entry into cell lines expressing low to moderate amounts of membrane associated DPP4 but is compatible with robust entry into cells expressing high levels of DPP4 at their surface. This resistance mechanism is related to that reported for HIV, where primary isolates exhibited markedly greater resistance to soluble CD4 (sCD4) than laboratory-adapted viruses (45, 46). The increased resistance was attributed to reduced sCD4 binding and diminished sCD4-induced shedding of gp120, the surface subunit of the viral envelope protein (45). Further, the resistance mechanism is similar to that described for SARS-CoV-2 -passaging of the virus in the presence of a dimeric ACE2-Fc protein resulted in acquisition of mutation Y489H in the viral S protein, which reduced binding to soluble ACE2 (47). Thus, SARS-CoV-2 and MERS-CoV can acquire resistance against soluble receptors by acquiring mutations that decrease receptor affinity and employing receptor decoys which bind the S protein with increased affinity and/or avidity can suppress resistance development (40, 48, 49).

It remains unknown whether authentic MERS-CoV would acquire resistance to sDPP4 through mutations at L507. However, we note that MERS-CoV sequences carrying L507F (n = 1, GenBank ID: QLD98042.1 (18)), L507R (n = 1, GenBank ID: ALW82691.1 (17)), or L507P (n = 2, GenBank IDs: AU54496.1 and AU54518.1 (17)) were detected in patients, suggesting that some of the mutations found in the present study that confer sDPP4 resistance might be compatible with viral spread in patients. Expression of membrane-associated DPP4 in the respiratory tract is thought to be highest in bronchus and lung while expression in the nasopharynx is comparatively low (50–52), which may partially account for the low human-human transmissibility of MERS-CoV. One can therefore speculate that mutations of residue L507 that reduce DPP4 binding might be compatible with some level of viral spread and disease pathogenesis in the lower respiratory tract while infection of the upper respiratory tract and transmission might be compromised.

Our findings await confirmation with authentic MERS-CoV and primary cells of the respiratory epithelium. Nevertheless, the results obtained with surrogate systems suggest that MERS-CoV can reduce DPP4 affinity in order to avoid inhibition by sDPP4 without losing its capacity to infect cells the lower respiratory tract. The development of sDPP4 as antiviral would thus benefit from DPP4 variants with augmented spike affinity,

## MATERIAL AND METHODS

### Expression plasmids

We used previously described expression plasmids encoding MERS-S from the human betacoronavirus 2c EMC/2012 isolate (GenBank: AFS88936.1)(53, 54), designated WT in the present study, the Korea outbreak–associated MERS-S I529T variant(16), DsRed(55), VSV-G (56), human DPP4(57), and soluble human DPP4 (14). Expression plasmids for the coding sequences of VSV nucleoprotein (VSV-N), phosphoprotein (VSV-P), and RNA-dependent RNA-polymerase (VSV-L) have also been described previously (58). To generate replication-competent, chimeric VSV expressing the MERS-CoV EMC spike protein (GenBank: AFS88936.1; VSV-MERS-S), a plasmid-encoded VSV anti-genome containing an additional transcription unit for eGFP between the VSV-G and VSV-L open reading frames (PMID: 29216247), was modified by replacing the native VSV-G gene by the sequence encoding the MERS-S EMC using engineered restriction sites. Mutant MERS-S expression constructs carrying L507H, L507I, L507F, L507R, L507P, T533P, L697F, L507H+L697F, or L507I+T533P+L697F substitutions were generated by overlap extension PCR using the MERS-S EMC plasmid as the template. For immunoblot-based detection, corresponding C-terminal V5-tagged versions were generated for each S protein. All plasmids were verified by automated Sanger sequencing (Microsynth SeqLab).

### Cell culture

Cell culture experiments were performed using Vero (African green monkey kidney; female; ATCC CRL-1586, RRID: CVCL_0603; kindly provided by Andrea Maisner), Huh-7 (human liver; male; JCRB JCRB0403, RRID: CVCL_0336; kindly provided by Thomas Pietschmann), 293T (human kidney; female; DSMZ ACC-635, RRID: CVCL_0063), and Caco-2 cells (human colon; male; ATCC HTB-37, RRID: CVCL_0025). All cell lines were propagated at 37°C in a humidified incubator containing 5% CO₂. Vero, Huh-7, and 293T cells were maintained in DMEM (PAN-Biotech) supplemented with 10% fetal bovine serum (FBS; Biochrom) and penicillin-streptomycin (PAN-Biotech; 100 U/ml penicillin and 0.1 mg/ml streptomycin). In contrast, Caco-2 cells were grown in MEM (Thermo Fisher Scientific) containing 10% FBS, 1% penicillin-streptomycin, 1% non-essential amino acids (PAA), and 1 mM sodium pyruvate (PAN-Biotech). Cell line identity was confirmed by short tandem repeat profiling and by amplification and sequencing of a cytochrome c oxidase gene fragment, complemented by microscopic inspection and/or evaluation of expected growth behavior. Mycoplasma testing was carried out routinely. For plasmid expression, 293T cells were transfected using calcium phosphate precipitation.

### Production of human sDPP4

Soluble human DPP4 fused to the Fc region of human IgG (sDPP4-Fc) was produced in 293T cells essentially as described previously (14). Briefly, 293T cells were transfected with an expression plasmid encoding sDPP4-Fc, supernatants were collected on days 4 and 5 posttransfection and clarified by centrifugation (2,000 × *g*, 10 min, 4°C). To maximize recovery, residual protein was additionally released from the remaining cells by two freeze–thaw cycles and combined with the clarified culture supernatants. The material was concentrated using 50-kDa MWCO centrifugal concentrators and purified using Protein A/G spin columns. Purified sDPP4 was further concentrated, aliquoted, and stored at −80°C until use.

### Viruses and infection

The replication-competent VSV-MERS-S reporter virus (described above) was rescued according to previously established protocols (58, 59). To evaluate infection dynamics, viral spread in the presence or absence of sDPP4 was subsequently monitored using this eGFP-expressing chimera.

### Verification of VSV-MERS-S integrity

The integrity of VSV-MERS-S stocks was assessed by sequencing of the spike gene. Viral RNA was purified using the QIAamp Viral RNA Mini Kit (Qiagen) and converted to cDNA with SuperScript III and random hexamers (Thermo Fisher Scientific). Spike-specific fragments were generated by PCR and nested PCR using Phusion polymerase, resolved on agarose gels, excised and purified (Macherey & Nagel), and subjected to automated Sanger sequencing (SeqLab)

### In vitro evolution under sDPP4 selection pressure

Virus stocks were titrated on Caco-2 cells by focus-forming assay (eGFP-based) as described (29) to determine focus-forming units (FFU) and calculate the multiplicity of infection (MOI) for initial infections. For in vitro evolution experiments, Caco-2 cells seeded in 24-well plates were infected with VSV-MERS-S at an MOI of 0.1 for 1 h at 37°C (5% CO₂). After adsorption, cells were washed with PBS and incubated with medium containing serial dilutions of in-house–produced sDPP4 (final dilutions: 1:10, 1:40, 1:160, 1:640, 1:2,560, and 1:10,240). Infection and spread were monitored by eGFP fluorescence microscopy. For each passage, supernatants were collected from the well containing the highest sDPP4 concentration that still yielded >90% eGFP-positive cells (typically 24–48 h postinfection), clarified by centrifugation to remove debris, aliquoted, and stored at −80°C. Subsequent passages were initiated by inoculating fresh Caco-2 cells with a 1:2,000 dilution of the previous passage supernatant under the same selection scheme. Supernatants from passages 5 and 10 were subjected to downstream phenotypic and genotypic analyses.

### Production of pseudotyped particles

VSV pseudotype particles were generated using a replication-deficient VSV backbone lacking the VSV-G gene and carrying dual reporters (eGFP and firefly luciferase), VSV*ΔG-FLuc (kindly provided by Gert Zimmer) (60), essentially as described previously (55). Particles were produced with either the indicated MERS-S proteins, VSV-G (positive control), or no viral glycoprotein (negative control). For the latter, producer cells were transfected with a DsRed expression plasmid to match transfection conditions. Briefly, 293T cells were transfected with plasmids encoding the respective glycoproteins. At 24 h posttransfection, cells were inoculated with VSV*ΔG-FLuc for 1 h at 37°C. The virus-containing inoculum was then removed, and the cells were washed with PBS. Fresh DMEM was added for virus production; to inactivate residual input virus carrying VSV-G, the medium was supplemented with anti-VSV-G antibody (I1 hybridoma supernatant; ATCC CRL-2700), except for cells transfected with the VSV-G expression plasmid, for which no neutralizing antibody was added. Producer cell supernatants containing pseudotyped particles were collected after 16–18 h, cleared by centrifugation to remove cell debris (4,000 × g, 10 min), aliquoted, and stored at −80°C until use.

### Immunoblot

To assess incorporation of MERS-S into VSV pseudotype particles, pseudoviruses carrying WT or the indicated mutant spikes (all equipped with a C-terminal V5 tag) were first concentrated by ultracentrifugation through a sucrose cushion (20% [wt/vol] sucrose in PBS; 16,000 rpm, 90 min, 4°C). Pelleted particles were resuspended and lysed in an equal volume of 2× SDS sample buffer (0.03 M Tris-HCl, 10% glycerol, 2% SDS, 5% β-mercaptoethanol, 0.2% bromophenol blue, 1 mM EDTA), denatured at 96°C for 10 min, and separated by SDS-polyacrylamide gel electrophoresis (SDS-PAGE). Proteins were transferred to nitrocellulose membranes (Hartenstein). Membranes were blocked for 30 min in PBS-T (PBS containing 0.05% Tween-20) supplemented with 5% skim milk and then incubated at 4°C overnight with antibodies against V5 (1:2,000; Proteintech; 14440-1-AP) or anti-VSV-M [23H12] (1:1,000; Kerafast; EB0011). After three washes with PBS-T, membranes were incubated for 1 h at room temperature with HRP-coupled secondary antibodies (anti-rabbit and anti-mouse; 1:2,000; Dianova; 115-035-045 and 115-035-003) diluted in 5% milk/PBS-T. Following three additional PBS-T washes, chemiluminescent detection was performed using Western blotting substrate (Cyanagen; XLS0142-0250). Signals were recorded on an Azure 600 imaging system and quantified with AzureSpot Pro software (Azure Biosystems). In addition, the same 293T producer cells used for pseudotype generation were harvested to assess S protein expression in whole-cell lysates. Cells were washed once with PBS and lysed directly in 2× SDS sample buffer, followed by heat denaturation at 96°C for 10–15 min. Lysates were analyzed by immunoblotting using an anti-V5 antibody (1:2,000; Proteintech; 14440-1-AP) for S expression, and GAPDH (1:2,000; Proteintech; 60004-1-Ig) served as a loading control. For quantification, S protein signals (S0 and S2) were normalized to the corresponding VSV-M or GAPDH signal and subsequently expressed relative to WT S protein (set to 1).

To assess cellular DPP4 expression, 293T, Vero, Huh-7, and Caco-2 cells were seeded in 6-well plates one day prior to harvest (2 × 10^5 cells/ml). The next day, medium was aspirated and cells were washed once with PBS. Cells were lysed directly in 2× SDS sample buffer for 10–15 min at room temperature, transferred to microcentrifuge tubes, and heat-denatured at 96°C for 15 min. Lysates were separated by SDS-PAGE and analyzed by immunoblotting as described above. DPP4 was detected using anti-DPP4 antibody (1:500; R&D Systems; AF1180). GAPDH was used as a loading control and detected with anti-GAPDH antibody (1:2000; Proteintech; 60004-1-Ig). HRP-conjugated anti-goat (1:2,000; Dianova; 705-035-003) and anti-mouse (1:2,000; Dianova; 115-035-003) secondary antibodies were used for detection.

### Spike protein–mediated entry and inhibition assays

To compare cell susceptibility and entry efficiency, target cells were plated in 96-well format and infected with identical volumes of VSV pseudotypes bearing the indicated MERS-CoV S proteins. Particles pseudotyped with VSV-G served as a positive control, while particles produced in the absence of a viral glycoprotein were included as a negative control. Following incubation for 16–20 h, transduction was quantified by measuring virus-encoded firefly luciferase activity in cell lysates. Briefly, culture medium was removed and cells were lysed for 30 min at room temperature in PBS containing 0.5% Tergitol 15-S-9 (Carl Roth; 50 µl per well). Lysates were transferred to white 96-well plates, combined with luciferase substrate (Beetle-Juice; PJK), and luminescence was recorded using a Hidex Sense plate luminometer (Hidex). For inhibition and neutralization experiments, pseudotyped particles were preincubated with entry inhibitors prior to inoculation. For soluble receptor blocking, virus preparations were mixed 1:1 with serial dilutions of sDPP4 (final dilutions: 1:10, 1:40, 1:160, 1:640, 1:2,560, and 1:10,240) and incubated at 37°C for 30–60 min. Particles incubated with medium alone served as the reference control. Luciferase signals were measured at 16–20 h postinoculation as described above.

### DPP4 binding efficiency and S protein surface expression assay

For analysis of spike surface expression and DPP4 binding by flow cytometry (FACS), 293T cells were plated in 6-well plates and transfected with expression plasmids encoding the respective S proteins or an empty vector (negative control). At 24 h posttransfection, the medium was replaced. At 48 h posttransfection, cells were collected for staining: cultures were washed, resuspended in PBS, and pelleted (600 × g, 5 min, 4°C). Cell pellets were washed once in PBS-B (PBS supplemented with 1% bovine serum albumin; BSA [Carl Roth]) and centrifuged again. For primary staining, cells were resuspended in PBS-B containing either anti–MERS-CoV S antibody to detect spike surface expression (1:1,000; Sino Biological; 40069-R723) or sDPP4-Fc to measure receptor binding (1:200; ACRO Biosystems; DP4-H5266) and incubated for 1 h at 4°C under gentle rotation (Rotospin, IKA). After washing, cells were incubated for an additional 1 h at 4°C with Alexa Fluor 488–conjugated secondary antibodies diluted in PBS-B: anti-rabbit IgG (1:200; Thermo Fisher Scientific; A-21206) for spike staining or anti-human IgG (1:200; Thermo Fisher Scientific; A-11013) for detection of bound sDPP4. Cells were then washed, resuspended in PBS-B, and analyzed on an ID7000 Spectral Cell Analyzer (Sony Biotechnology).

### DPP4 expression assay in different cell lines by flow cytometry

For flow-cytometric analysis of DPP4 surface expression, Vero, Huh-7, 293T, and Caco-2 cells were seeded in 6-well plates 2 days prior to staining, and culture medium was refreshed the next day. In parallel, 293T cells transfected with a human DPP4 expression plasmid were included as a positive control (medium change at 24 h posttransfection). On the day of analysis, cells were washed once with PBS and detached using trypsin–EDTA (0.05% trypsin/0.02% EDTA; PAN; P10-023100), Accutase (PAN; P10-21500), or citric saline buffer (KCl and sodium citrate in Milli-Q water) for 5–8 min at 37°C, followed by preparation of single-cell suspensions. Cells were pelleted (600 × g, 10 min, 4°C) and washed three times with PBS-B. Cells were then incubated with an anti-DPP4 primary antibody (Abcam; ab119346) for 60 min at 4°C, washed twice with PBS-B, and stained with Alexa Fluor 488–conjugated anti-mouse IgG (Thermo Fisher Scientific; A-21200) for 60 min at 4°C. After two additional washes with PBS-B, cells were resuspended in PBS-B and analyzed by flow cytometry as described above.

### Three-dimensional RBD models

Illustration of mutation L507I in the context of the MERS-CoV S protein structure was performed using the crystal structure PDB: 5X59 (61) as template. Visualization and coloring was done using the UCSF Chimera (version 1.17.3) software (62).

### Data analysis

Data analysis was conducted using Microsoft Excel (as part of the Microsoft Office software package, version 2016, Microsoft Corporation, Redmond, WA, USA) and GraphPad Prism version 10.6.1 (GraphPad Software, San Diego, CA, USA). Depending on the experiment, significance testing was carried out using either a one-way ANOVA followed by Dunnett’s posttest (particle incorporation, spike-driven entry, sDPP4-mediated inhibition in Vero cells, and sDPP4 binding assessed by flow cytometry), or two-way ANOVA with Dunnett’s posttest (sDPP4 inhibition in Caco-2 cells and mAb inhibition assays in Vero cells). The sDPP4 dilution factor that results in 50% inhibition (inhibitory dilution 50%, Log10 ID50) was calculated using a non-linear regression model. For negative Log10 ID50 values, values were manually set as 0. Only p-values of 0.05 or lower were considered statistically significant (p > 0.05, not significant [ns]; p ≤ 0.05, *; p ≤ 0.01, **; p ≤ 0.001, ***).

## Data availability

All data generated during this study are included in this article. GenBank accession numbers cited refer to sequences previously deposited by others and are publicly available through the National Center for Biotechnology Information (NCBI) database.

## Acknowledgements

We thank Roberto Cattaneo, Michael Farzan, Georg Herrler, Stephan Ludwig, Andrea Maisner, Vincent Munster, Thomas Pietschmann, and Gert Zimmer for providing reagents.

